# Spaces of phylogenetic diversity indices: combinatorial and geometric properties

**DOI:** 10.1101/2023.02.02.526891

**Authors:** Kerry Manson, Mike Steel

## Abstract

Biodiversity is a concept most naturally quantified and measured across sets of species. However, for some applications, such as prioritising species for conservation efforts, a species-by-species approach is desirable. Phylogenetic diversity indices are functions that apportion the total biodiversity value of a set of species across its constituent members. As such, they aim to measure each species’ individual contribution to, and embodiment of, the diversity present in that set. However, no clear definition exists that encompasses the diversity indices in current use. This paper presents conditions that define diversity indices arising from the phylogenetic diversity measure on rooted phylogenetic trees. In this context, the diversity index ‘score’ given to a species represents a measure of its unique and shared evolutionary history as displayed in the underlying phylogenetic tree. Our definition generalises the diversity index notion beyond the popular Fair Proportion and Equal-Splits indices. These particular indices may now be seen as two points in a convex space of possible diversity indices, for which the boundary conditions are determined by the underlying shape of each phylogenetic tree. We calculated the dimension of the convex space associated with each tree shape and described the extremal points.

## 1 Introduction

The evolutionary connections and relationships within sets of species are most commonly modelled by phylogenetic trees [1]. Applying quantitative measures to such trees has proven useful in understanding these relationships and for highlighting conservation priorities in the face of the current human-induced mass extinction of species [2]. A large number of these quantitative approaches have been developed (see [3] for an overview).

Phylogenetic diversity (PD) is a measure that aims to quantify the biodiversity exhibited by a set of species [4]. PD does so by using a weighted phylogenetic tree that exhibits the evolutionary relationships among species in the set, where the weights represent time elapsed or genetic change along the edges of the tree. In broad terms, a PD value for a phylogenetic tree is calculated by adding the weights of all the edges in that tree together. This sum can be seen to represent the total evolutionary history shared by the species at the leaves. However, in some contexts, it is useful to ask how much of that total can be attributed to each species on an individual basis. This perspective opens up the possibility of ranking species by their ‘distinctiveness’ [5] or ‘evolutionary isolation’ [6], or to quantify their ‘combination of unique and shared evolutionary heritage’ [7]. Doing so provides evidence for developing conservation priorities.

The usual means of arriving at such a ranking is by way of a (phylogenetic) diversity index, viewing the weights as edge ‘lengths’. These methods have been described as ‘distributing edge lengths among descending leaves’ [8], a characterisation that we shall use.^1^ The most commonly used diversity indices are the Fair Proportion (FP) index and the Equal-Splits (ES) index (described below). The former (under the name Evolutionary Distinctiveness) is used by the EDGE of Existence programme [11–13] to help rank species by their need for conservation assistance. A recent study by Palmer and Fischer [14] evaluated the effects of this implementation of Evolutionary Distinctiveness on conservation efforts.

Although these particular indices are built on clear principles, they are by no means the unique solutions to the problem of allocating the total PD value among the leaves. No general definition has yet appeared to encompass both the known diversity indices and further possibilities. In this paper, each particular allocation of the PD value is framed in terms of a set of coefficients that determine it. We then give a pair of conditions on these coefficients that satisfy some natural allocation requirements within a simple evolutionary model. These conditions define diversity index functions in a general sense. Some further constraints on the coefficients are derived as immediate consequences of our definition of a diversity index. We also show that, given a set of diversity indices, we can create further diversity indices by taking linear combinations of the original ones. This observation, in turn, leads to descriptions of convex spaces, within which all of the diversity indices for a given tree shape may be positioned. We show how the topology of the underlying tree (the ‘tree shape’) determines the dimension and boundaries of its associated diversity index space.

The FP and ES indices are thus recast as single points within these spaces. We describe the diversity indices that lie at the extreme points of the convex spaces, from which, by way of convex combination, every index in the space can be calculated. This provides a means of defining the full range of solutions to the allocation problem and the relationships among these possibilities. We illustrate these results by describing, in detail, the convex spaces associated with the three rooted binary tree shapes containing five leaves, and by including a case study of the phylogenetic tree of hominoids (great apes and gibbons).

With this method of determining every diversity index on a particular tree, we can investigate the properties of certain diversity indices and see whether or not these properties are unique to that index or if they are widely held. For example, we used our diversity index conditions to show that the FP index is the unique diversity index that obeys a certain ‘continuity’ condition on all tree shapes. This indicates that the FP index is the most appropriate to use with rooted trees that are not fully resolved. Another natural property, which we call ‘consistency’ and which is shared by both the FP and ES indices, is discussed. We show that given fixed edge lengths, every diversity index that satisfies our definition is able to be framed as an equivalent consistent one. This allows us to view indices as a process of re-weighting edge lengths, akin to a flow problem, and to describe the convex spaces of diversity indices in more detail. Our concluding remarks discuss other similar conditions and suggest families of diversity indices that may be of further interest.

## 2 Preliminaries

We begin by recalling some standard terminology of phylogenetic trees and then introduce some additional notions. Let *X* be a non-empty set of taxa (e.g. species), with |*X*| = *n*. A *rooted phylogenetic X-tree* is a rooted tree *T* = (*V, E*), where *X* is the set of leaves, and all edges are directed away from a distinguished root vertex *ρ*, and every non-leaf vertex has out-degree at least 2. We call these non-leaf vertices *interior* vertices. In addition, when |*X*| = 1, the tree consisting of a single vertex is a rooted phylogenetic *X*-tree. Ignoring the labelling of the leaves by elements of *X* gives us the *tree shape*. If all interior vertices of *T* have out-degree 2, we say that *T* is *binary*.

All edges drawn in this paper will be directed down the page. For the directed edge *e* = (*u, v*), we say that *u* is the *initial* vertex and that *v* is the *terminal* vertex. We also say that *u* is the *parent* of *v* and *v* is a *child* of *u*. An edge *e* from *E*(*T*) is *pendant* if its terminal vertex has out-degree zero. Otherwise, *e* is an *interior* edge. A subtree of *T* is *pendant* if it can be disconnected from *ρ* by deleting a single edge of *T* . We use *P*_*e*_ to denote the pendant subtree formed by deleting the edge *e*.

We say that a vertex *v* is *descended from* vertex *u* in *T* if *u* is distinct from *v* and there exists a directed path in *T* from *u* to *v*. We also say that an edge *e* is descended from the distinct edge *f* if the terminal vertex of *e* is descended from the terminal vertex of *f* . Edges descended from vertices are defined similarly, by reference to the terminal vertex of the edge involved. However, for vertices descended from edges we do not require the vertex and the terminal vertex of the edge to be distinct. A set *S* of vertices and/or edges may be said to be descended from an edge *e* (resp. vertex *v*) if each member of *S* is descended from *e* (resp. *v*). The *cluster* of all leaves descended from edge *e* is denoted as *c*_*T*_ (*e*).

Let *e* be an interior edge of *T* for which the terminal vertex *v* has outdegree *d*. We represent the *d* maximal pendant subtrees contained within *P*_*e*_ by *T*_1_(*e*), *T*_2_(*e*), …, *T*_*d*_(*e*). Depending on the context, it may be useful to denote these subtrees by *T*_1_(*v*), *T*_2_(*v*), *…, T*_*d*_(*v*), and allow this notation to extend to the case where *v* is the root vertex. Figure 1 illustrates this notation for an edge *e* where *d* = 2.

**Fig. 1:**
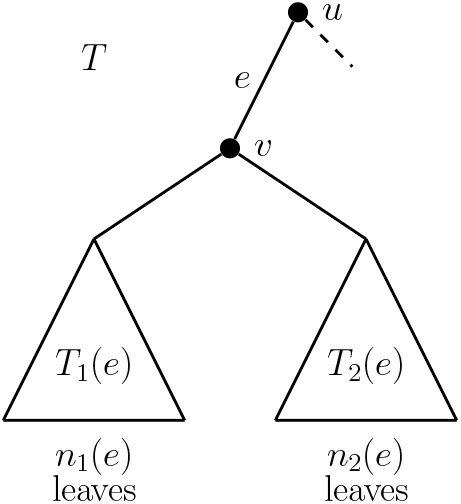
Schematic diagram showing the notation for maximal pendant subtrees descended from edge *e*. The vertex *v* has out-degree 2; however, in general, we allow larger out-degrees.

The edges of every rooted phylogenetic tree considered in this paper are positively weighted. We call these weights *edge lengths*. For a rooted phylogenetic *X*-tree *T*, let *ℓ* : *E*(*T*) → ℝ^*>*0^ be a function that assigns a positive real-valued length *ℓ* (*e*) to each edge *e* ∈ *E*(*T*). We refer to *ℓ* as an *edge length assignment function*. We make no further restrictions on the edge lengths other than positivity (i.e., no ultrametric condition is enforced). The *phylogenetic diversity* of *T* given edge length assignment function *ℓ*, denoted *PD*(*T, ℓ*), is defined as the sum of the edge lengths of *T* . That is:

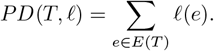

Two functions that we considered frequently in this paper were the Fair Proportion (FP) index [13, 15] and the Equal-Splits (ES) index [6, 15]. Let *P* (*T* ; *ρ, x*) be the path in *T* from the root vertex *ρ* to leaf vertex *x*. For each leaf *x* ∈ *X*, the *Fair Proportion* index score of *x* in *T* is given by:

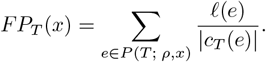

For each leaf *x* ∈ *X*, the *Equal-Splits* index score of *x* in *T* is given by:

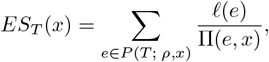

where Π(*e, x*) is the product of the out-degrees of the interior vertices appearing on the path from the terminal vertex of *e* to *x*, when *e* is an interior edge, and Π(*e, x*) = 1, when *e* is incident to *x*.

## 3 Diversity indices

In this section, we provide a set of conditions defining phylogenetic diversity indices in general, for rooted phylogenetic trees. Our definition is guided by a standard biological interpretation of rooted phylogenetic tree structures. For instance, the root vertex corresponds to the most recent common ancestor of the leaves (species) in its tree. Thus the path from the root to a leaf traces the evolutionary development of that leaf species from the common ancestor through to the present. Additionally, the length of each edge in a rooted phylogenetic tree is assumed to reflect the amount of evolutionary change that occurred along that edge. Moreover, evolutionary changes common to the cluster of species *c*_*T*_ (*e*) are explained by changes occurring somewhere along the edge *e*. As such, the process of speciation aligns with the tree shape of a rooted phylogenetic tree, where the interior vertices correspond to speciation events.

It is against this interpretation (and the observations above) that we test the conditions that define diversity indices. The broad goal of any phylogenetic diversity index is to take the overall PD score of a rooted phylogenetic tree and distribute this value among the species in a way that is compatible with the tree shape. We aim to present this distribution in a general form, but not to the extent that species are given negative values. A negative allocation of PD value to a species would be equivalent to saying that that species was taking away from the phylogenetic diversity of other species by continuing to survive.

We begin Section 3.1 by describing a more general class of functions, here called allocation functions, as well as the coefficient notation that will be used. The further conditions required of diversity indices are included in Section 3.2. Definition 2 encompasses the FP and ES indices defined above and allows for the description of further diversity indices. The distinctions between individual diversity indices should reflect different assumptions about how species exhibit ancestral developments while respecting the observations noted above. We conclude this section with some results on the index coefficients that arise immediately from this definition.

### 3.1 Allocation functions

#### Definition 1

Let *T* = (*V, E*) be a rooted phylogenetic *X*-tree with edge length assignment function *ℓ*. An *allocation function φ* _*ℓ*_ : *X* → ℝ^≥0^ is a real-valued function on the set of leaves of *T* that satisfies the following equation:

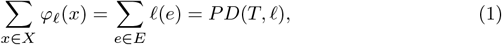

and moreover it may be expressed as *φ*_*ℓ*_ (*x*) = Σ_*e*∈*E*_ *γ*(*x, e*) *ℓ* (*e*), for every edge length assignment function *ℓ*, where all of the *coefficients γ*(*x, e*) are non-negative.

We call the value *φ*_*ℓ*_ (*x*) the *φ-score* of leaf *x* (given the length assignment *ℓ*). Our aim is to define diversity indices that not only deliver an ordinal ranking of species according to their contribution to PD, but also to measure this contribution. With this in mind, it is sensible to require allocation functions (and, consequently, diversity indices) to partition the total PD value of a rooted phylogenetic tree among its leaf species. This idea is expressed by Equation (1) and is reinforced by requiring each allocation function to be expressed in terms of coefficients *γ*(*x, e*) that do not depend on the particular edge length assignment function *ℓ*. Edge lengths will be used to calculate the value of individual allocation function scores but do not themselves impact the method of allocation. Because of this independence, the length subscript on *φ*_*ℓ*_will be omitted when *ℓ* is clear from the context.

The final condition of Definition 1, that all of the coefficients *γ*(*x, e*) are non-negative, is worth considering further. Without it, the class of allocation functions would contain many biologically unreasonable functions. One such unreasonable allocation can be described on the small rooted phylogenetic tree with exactly two leaves *x* and *y*, and two edges *a* and *b*. On this tree, consider the function *σ* : {*x, y*} → ℝ, defined as follows:

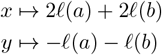

Then *σ* indeed satisfies (1), as *σ*(*x*) + *σ*(*y*) = *ℓ* (*a*) + *ℓ* (*b*). However, claiming that *σ*(*y*) represented the evolutionary history of *y* would be hard to justify. Negative coefficients and scores do not fit our intended model, where diversity indices act as a measure of (necessarily positive) evolutionary history; hence the final stipulation in Definition 1. The next result shows that, similar to PD, allocation functions are *linear* in the following sense.

#### Proposition 1

*Let T* = (*V, E*) *be a rooted phylogenetic X-tree. Furthermore, let φ be an allocation function that may be written in the form φ* _*ℓ*_ (*x*) = Σ_*e* _∈_*E*_ *γ*(*x, e*) *ℓ* (*e*) *for every edge length assignment function ℓ. Suppose that s, t* ∈ ℝ *and that ℓ, ℓ*_1_ *and ℓ*_2_ *are edge length assignment functions such that ℓ* (*e*) = *sℓ*_1_(*e*) + *tℓ*_2_(*e*) *for every edge e* ∈ *E. Then* 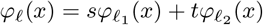 *for all leaves x* ∈ *X*.

*Proof* Let *s, t* ∈ ℝ and suppose that *ℓ, ℓ*_1_ and *ℓ*_2_ are edge length assignment functions such that *ℓ* (*e*) = *sℓ*_1_(*e*) + *tℓ*_2_(*e*) for every *e* ∈ *E*. Then:

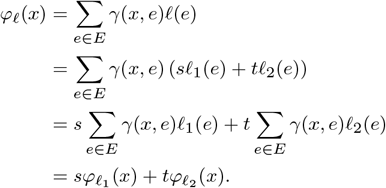

□

An allocation function on a phylogenetic *X*-tree *T* = (*V, E*) may be determined by a rule or formula, or, if needed, can be completely described by listing the |*X*| × |*E* |coefficients *γ*(*x, e*). Note that the FP and ES indices satisfy Definition 1, and hence are allocation functions. For these indices, the coefficients may be read directly from their definitions. That is, they can be read as 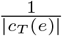 and 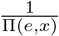, respectively, when *x* ∈ *c*_*T*_ (*e*), noting that for both indices *γ*(*x, e*) = 0 whenever *x* is not descended from *e*. In contrast, we can define the coefficients directly. Let |*X*| = *n*, and take 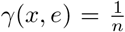 for all *x* ∈ *X* and all *e* ∈ *E*. Then *φ* (*x*) = Σ_*e* _∈_*E*_ *γ*(*x, e*) *ℓ* (*e*) is an allocation function that shares the total PD value of *T* uniformly among the *n* leaves.

For some tree shapes, two allocation functions with different descriptions or formulae for calculating coefficients may nevertheless ultimately produce the same scores. We say that two allocation functions *φ* and *ψ coincide* if *φ* _*ℓ*_ (*x*) = *ψ* _*ℓ*_ (*x*) for every *x* ∈ *X* and every edge length assignment function *ℓ*. This concept has been used to understand the similarities and differences among some diversity indices. For instance, Wicke and Steel [7] characterised the rooted tree shapes for which FP and ES coincide.

### 3.2 Diversity indices

The ‘uniform’ allocation function above, where each coefficient is 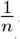, is an allocation function that ignores the phylogenetic structure entirely. It is thus an allocation function that does not allow for relative or differing contributions to biodiversity. In contrast, diversity indices are a subclass of allocation functions that does account for the rooted phylogenetic tree structure.

We intend for *γ*(*x, e*) to represent the proportion of evolutionary history that arises along edge *e* that is currently embodied by species *x*. This provides our first constraint, namely that the evolutionary history arising along edge *e* should be allocated exclusively to species descended from *e*.

Additionally, the set of coefficients for an edge should be entirely determined by the shape of its descendent tree structure, not by any ancestral or otherwise unconnected parts of the tree. In other words, the same pattern of descent should lead to the same pattern of coefficients. Furthermore, the particular labelling of the leaves should be inconsequential for the calculation of a diversity index. Definition 2 (below) restricts the class of allocation functions to those that obey these minimal criteria. In order to describe this definition, we first describe two further notions.

A *symmetry* of *T* is a permutation of the vertices that maintains precisely those relationships of descent found in *T* . Suppose that *T* and *T′* are two rooted phylogenetic trees with the same tree shape. Let *π* be a bijective map from the vertices of *T* to the vertices of *T′*, such that *v* is descended from *u* if and only if *π*(*v*) is descended from *π*(*u*). Then *v* and *π*(*v*) are said to be in *corresponding positions* of that tree shape. Moreover, recall that *P*_*e*_ denotes the pendant subtree formed by deleting the edge *e*.

#### Definition 2

Let *T* = (*V, E*) be a rooted phylogenetic *X*-tree. A *diversity index* (on *T*) is an allocation function *φ*_*ℓ*_: *X* → ℝ^*>*0^ given by *φ* _*ℓ*_ (*x*) = Σ_*e* _∈_*E*_ *γ*(*x, e*) *ℓ* (*e*) for every edge length assignment function *ℓ*, that additionally satisfies the conditions (DI_1_) and (DI_2_) below:

• (DI_1_) *Descent condition: γ*(*x, e*) = 0 if *x* is not descended from *e*.

• (DI_2_) *Neutrality condition:* The coefficients *γ*(*x, e*) are a function of the tree shape of *P*_*e*_. Moreover, suppose that *P*_*e*_ and *P*_*f*_ are pendant subtrees of *T* with the same tree shape. If leaves *x* in *P*_*e*_ and *y* in *P*_*f*_ appear in corresponding positions in their respective subtrees, then *γ*(*x, e*) = *γ*(*y, f*).

Consequently, to satisfy the neutrality condition (DI_2_), any diversity index on the tree in Figure 2 is required to satisfy all of the following coefficient equalities: *γ*(*x*_1_, *a*) = *γ*(*x*_2_, *a*), *γ*(*x*_5_, *b*) = *γ*(*x*_6_, *b*) = *γ*(*x*_1_, *c*) = *γ*(*x*_2_, *c*), *γ*(*x*_7_, *b*) = *γ*(*x*_3_, *c*) and *γ*(*x*_5_, *d*) = *γ*(*x*_6_, *d*) = *γ*(*x*_1_, *e*) = *γ*(*x*_2_, *e*).

**Fig. 2:**
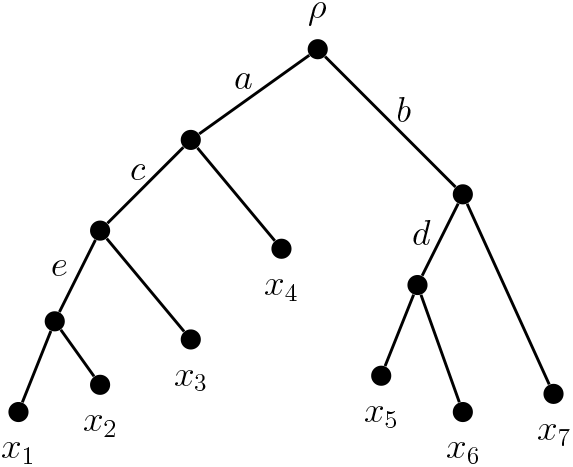
Any diversity index on the above tree must allocate the length of edge *c* exclusively among its descendent cluster: {*x*_1_, *x*_2_, *x*_3_ }. The pattern of this allocation must be matched by the allocation of the length of edge *b* among *b*’s own descendent cluster, namely {*x*_5_, *x*_6_, *x*_7_ }. This is because the maximal pendant subtrees below *b* and *c* have the same tree shape.

We now discuss some immediate consequences of our definition. Proposition 3 discusses some constraints on the allocation coefficients. Proposition 4 reframes the neutrality condition in terms of symmetries, and Proposition 5 gives bounds on the index scores of a general leaf. Wicke [16] showed that for any function *φ* _*ℓ*_ (*x*) = Σ_*e* _∈_*E*_ *γ*(*x, e*) *ℓ* (*e*) that satisfies Equation (1), the coefficients associated with each edge sum to one. Lemma 2 reframes this result slightly so it applies to allocation functions.

#### Lemma 2

*Let T* = (*V, E*) *be a rooted phylogenetic X-tree with edge length assignment £*.*Suppose that γ*(*x, e*) ≥ 0 *for all x* ∈ *X and e* ∈ *E. Then the function φ* (*x*) = Σ_*e* _∈_*E*_ *γ*(*x, e*) *ℓ* (*e*) *is an allocation function if and only if* Σ_*x*∈*X*_ *γ*(*x, e*) = 1 *for every edge e* ∈ *E*.

#### Proposition 3

*Let T* = (*V, E*) *be a rooted phylogenetic X-tree with edge length assignment function ℓ. Let φ* (*x*) = Σ_*e* _∈_*E*_ *γ*(*x, e*) *ℓ* (*e*) *be a diversity index. Then*

(i) *γ*(*x, e*) ≤ 1 *for all x* ∈ *X and e* ∈ *E*,

(ii) 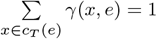, *and*

(iii) *if e is a pendant edge, then γ*(*x, e*) = 1 *when leaf x*′ *is incident with e, and γ*(*x, e*) = 0 *otherwise*.

*Proof* Suppose that *γ*(*x*′, *e*) *>* 1 for some edge *e* and leaf *x*′. From the definition of allocation functions, all other coefficients of *φ* are non-negative, so Σ_*e* _∈_*E*_ *γ*(*x, e*) *>* 1. However this contradicts Lemma 2 and hence *γ*(*x, e*) ≤ 1 for all *x* ∈ *X* and *e* ∈ *E*. Next, for a specified edge *e*, we can split the leaves into two sets: the set of leaves descended from *e*, denoted *c*_*T*_ (*e*), and the rest: *X* \ *c*_*T*_ (*e*). Then, starting from the Lemma 2 result:

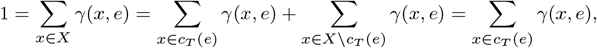

where the last equality follows from the descent condition (DI_1_). Lastly, let *e* be a pendant edge. Suppose *x* ′ ∈ *X* is incident with *e*, giving *c*_*T*_ (*e*) = {*x* ′}. Thus by Part (ii) 1 = Σ_*e* _∈_*E*_ *γ*(*x, e*) = *γ*(*x* ′, *e*). Now suppose that *x* ′ is not incident with *e*. Then *x* ′ is not descended from *e*, so by using (DI_1_) we conclude that *γ*(*x* ′, *e*) = 0.

□

#### Proposition 4

*Let T* = (*V, E*) *be a rooted phylogenetic X-tree with edge length assignment function ℓ. Let φ* (*x*) = Σ_*e* _∈_*E*_ *γ*(*x, e*) *ℓ* (*e*) *be a diversity index. For distinct leaves x, x* ′, *both descended from e, if there is a symmetry in T that swaps x for x* ′ *then γ*(*x, e*) = *γ*(*x* ′, *e*).

*Proof* Let *T* ′ be the resulting tree after applying the symmetry to *T* that swaps *x* for *x* ′. Then *P*_*e*_ and 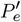 (the pendant subtrees created by deleting edge *e* in *T* and *T* ′, respectively) have the same tree shape, and leaf *x* in *P*_*e*_ corresponds to leaf *x* ′ in 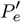. Hence, by the neutrality condition (DI_2_) *γ*(*x, e*) = *γ*(*x* ′, *e*). □

#### Proposition 5

*Let T* = (*V, E*) *be a rooted phylogenetic X-tree with edge length assignment function ℓ. Let e*_*x*_ *be the pendant edge incident with leaf x* ∈ *X, and let P* (*T* ; *ρ, x*) *be the path in T from the root vertex ρ to x. Let σ*_*x*_(*e*) *be the number of leaf vertices in c*_*T*_ (*e*) *that appear in a corresponding position to x (including x). If φ is a diversity index on T, then* 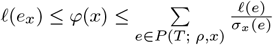 *for all x* ∈ *X*.

*Proof* Let *x* ∈ *X* and *φ* (*x*) = Σ_*e* _∈_*E*_ *γ*(*x, e*) *ℓ* (*e*). First suppose that *γ*(*x, e*) = 0 for all *e* ∈ *E* \ {*e*_*x*_}. This is the minimal possible choice, as Proposition 3(iii) ensures that *γ*(*x, e*_*x*_) = 1. In this case, 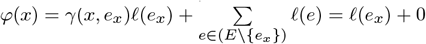.

Next, for some edge *e* ∈ *E*, suppose that *γ*(*x, e*) is non-zero. By (DI_1_), a nonzero coefficient means that *x* is descended from *e* (equivalently, *e* ∈ *P* (*T* ; *ρ, x*)). The coefficients of the *σ*_*x*_(*e*) leaves in corresponding positions to *x* must all have coefficients equal to *γ*(*x, e*). We require 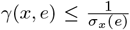 in order fofr the coefficients associated with *e* to sum to at most 1 and not contravene Proposition 3(ii). Thus 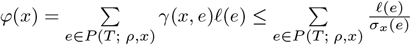. Therefore, for all *x* ∈ *X*,

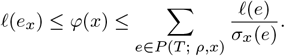

□

## 4 Continuity

In [17][p. 140], the following property of the FP index was noted:

“The index *ψ* = *FP* satisfies the following continuity condition: If *e* is an interior edge of a phylogenetic tree and *T/e* is the tree obtained from *T* by collapsing edge *e*, then 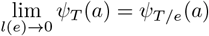.’’

As an illustration of how Definition 2 can be used to investigate the properties of diversity indices, we show that FP is the unique diversity index that satisfies this continuity condition on every tree. We use the same notation of *T/e* to denote the tree obtained from *T* by collapsing edge *e*, and refer to this property as the *diversity index continuity property*.

### Theorem 6

*Let ψ be a diversity index. Then* lim_. *ℓ* (*e*)→0_ *ψ*_*T*_ (*x*) = *ψ*_*T/e*_(*x*) *for every rooted phylogenetic X-tree T* = (*V, E*), *every x* ∈ *X and every interior edge e* ∈ *E*(*T*) *if and only if ψ is the Fair Proportion index*.

*Proof* Suppose that *ψ* is a diversity index that satisfies the diversity index continuity property on every rooted phylogenetic tree. Let *T* = (*V, E*) be a rooted phylogenetic *X*-tree with edge length assignment function *ℓ*. Since *ψ* is a diversity index, we have *ψ*_*T*_ (*x*) = Σ_*e* _∈_*E*_ *γ*(*x, e*) *ℓ* (*e*), for coefficients *γ*(*x, e*) that satisfy (DI_1_) and (DI_2_). The index may be defined by its set of coefficients. Moreover, we need to specify only those coefficients that are not already determined by the definition of a diversity index.

Let |*X*| = *n* and *E*(*T*) = {*e*_1_, *e*_2_, *…, e*_2*n*−2_}, where the pendant edges of *T* are indexed by {1, …, *n*} and the interior edges of *T* are indexed by {*n* + 1,, 2*n* − 2}. Let *e*_*i*_ be an interior edge of *T* and let *n*_*i*_ = |*c*_*T*_ (*e*_*i*_)|, the number of leaves of *T* descended from *e*_*i*_. First, assume that no interior edge of *T* is descended from *e*_*i*_. That is, only pendant edges appear below *e*_*i*_. By the neutrality condition (DI_2_) and Proposition 3(ii), if *x* is descended from *e*_*i*_, then 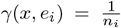. Otherwise, *γ*(*x, e*_*i*_) = 0.

Now assume that *e*_*j*_ is an interior edge descended from *e*_*i*_. By the diversity index continuity property:

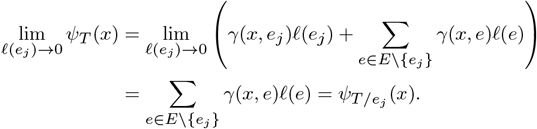

Note that the process of contracting an interior edge does not alter the coefficients *γ*(*x, e*) of *ψ*. Nor does it alter the number of leaves descended from any other edge. Let *D*(*e*_*i*_) be the set of interior edges descended from *e*_*i*_. We contract each edge from *D*(*e*_*i*_) in turn. By repeated use of the diversity index continuity property,

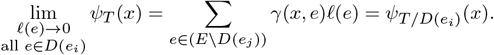

In *T/D*(*e*_*i*_), no interior edge is descended from *e*_*i*_. Hence, we again have 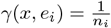 if *x* is a descendant of *e*_*i*_. Since the coefficients are unaffected by the contraction process, 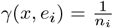 in *T* as well. By repeating this process for each interior edge *e*_*n*+1_, …, *e*_2*n*−2_, we obtain every coefficient *γ*(*x, e*) of each edge *e* ∈ *E*(*T*). This argument shows that in every case 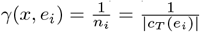, which is exactly the set of coefficients that defines the Fair Proportion index on *T* . Therefore, *ψ* is the FP index.

Conversely, suppose that *ψ* is the FP index. Then the coefficients *γ*(*x, e*) depend only on the number of leaves descended from edge *e*, and not on the particular structure of the phylogenetic tree below *e*. The contraction of any interior edge *e*_*i*_ does not reduce the number of leaf vertices descended from any other edge *e*. Hence the coefficients *γ*(*x, e*) are not altered when an interior edge distinct from *e* itself is contracted, and the diversity index continuity property holds. □

FP may therefore be considered the most appropriate diversity index to apply to rooted phylogenetic trees that are not fully resolved, as later refinements of the tree will not strongly impact the initial FP index scores. We note also that FP is special amongst diversity indices for a quite different reason — it is precisely the Shapley value for allocating the total phylogenetic diversity of a tree amongst the leaves [18].

## 5 Spaces of diversity indices

Each rooted phylogenetic tree has a restricted collection of diversity indices that may be applied to it. In this section, we discuss such collections of diversity indices. We show that these indices, when expressed as vectors of index scores, lie inside convex spaces determined by the structure of the associated tree. Examples of these spaces are presented for small rooted phylogenetic trees.

Let *T* = (*V, E*) be a rooted phylogenetic *X*-tree with *X* = {*x*_1_, …, *x*_*n*_} and *E* = {*e*_1_, …, *e*_*m*_}. Suppose that *φ* is a diversity index on *T* given by *φ* (*x*) = Σ_*e* _∈_*E*_ *γ*(*x, e*) *ℓ* (*e*) for the edge length assignment function *ℓ*. We place the coefficients *γ*(*x, e*) in an *n* × *m* matrix *A*_*φ*_ = (*a*_*ij*_), where *a*_*ij*_ = *γ*(*x*_*i*_, *e*_*j*_). Consider the tree in Figure 3. Its matrix of diversity index coefficients fits the pattern shown in the same figure, where the value of 0 ≤ *α* ≤ 1 is determined by the particular index. For example, FP uses 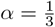 here and ES uses 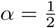. We now use the matrix of coefficients to determine the *ϕ* index scores of each leaf and subsequently form a space containing all the possible scores. The next two definitions introduce these concepts.

### Definition 3

Let *T* = (*V, E*) be a rooted phylogenetic *X*-tree with the leaf set *X* = {*x*_1_, …, *x*_*n*_} and edges *E* = {*e*_1_,*…, e*_*m*_}. Let *φ* be a diversity index on *T*, for which *A*_*φ*_ is the associated matrix of coefficients. The *index score vector* for *φ* is the *n*-component vector ***v***_***φ***_ = *A*_*φ*_***l***, where ***l*** is the vector of edge lengths: [*ℓ* (*e*_1_), *ℓ* (*e*_2_),…, *ℓ* (*e*_*m*_)]^*T*^ .

### Definition 4

Let *T* = (*V, E*) be a rooted phylogenetic tree. The *space of index score vectors* on *T*, denoted *S*(*T, ℓ*), contains all the index score vectors of *T* .

Note that *S*(*T, ℓ*) is not a vector space, since **0** is not in *S*(*T, ℓ*) for any edge length assignment function *ℓ*. This is because the index score of any leaf is always at least the (strictly positive) length of its incident pendant edge. However, *S*(*T, ℓ*) is convex, as we now show.

### Proposition 7

*Let T* = (*V, E*) *be a rooted phylogenetic X-tree. For any edge length assignment function ℓ, the space of index score vectors S*(*T, ℓ is convex*.

*Proof* Let *X* = {*x*_1_, …, *x*_*n*_} and *E* = {*e*_1_, …, *e*_*m*_}, and supposethat *φ* and *ψ* are diversity indices on *T*, where *φ* (*x*) = Σ_*e* _∈_*E*_ *γ*(*x, e*) *ℓ* (*e*) and *ψ*(*x*) = Σ_*e* _∈_*E*_ *γ* ′ (*x, e*) *ℓ* (*e*). Let *Aφ* = (*a*_*ij*_) = (*γ*(*x*_*i*_, *e*_*j*_)) and *B*_*ψ*_ = (*b*_*ij*_) = (*γ* ′ (*x*_*i*_, *e*_*j*_)) be the respective *n* × *m* coefficient matrices of *φ* and *ψ*. Finally, let ***u*** = *A*_*φ*_***l*** and ***v*** = *B*_*ψ*_ ***l*** be the respective vectors of index scores for *φ* and *ψ*. That is, ***u, v*** ∈ *S*(*T, ℓ*).

We first prove that the linear combination ***w*** = *t****u*** + (1 − *t*)***v*** is an allocation function for all real values of *t* ∈ [0, 1]. Let *δ*(*x*_*i*_, *e*_*j*_) = *tγ*(*x*_*i*_, *e*_*j*_) + (1 − *t*)*γ* ′ (*x*_*i*_, *e*_*j*_) for all 1 ≤ *i* ≤ *n* and 1 ≤ *j* ≤ *m*. We then have:

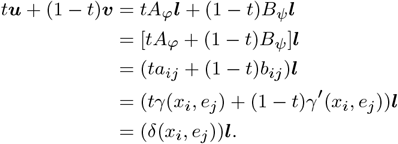

We form a new matrix *C* = (*c*_*ij*_) = (*δ*(*x*_*i*_, *e*_*j*_)) from these values. The coefficients *γ*(*x*_*i*_, *e*_*j*_) and *γ* ′ (*x*_*i*_, *e*_*j*_) are non-negative for all 1 ≤ *i* ≤ *n* and 1 ≤ *j* ≤ *m*, and for every *t* ∈ [0, 1] we have *t* ≥ 0 and (1 − *t*) ≥ 0. Thus the coefficients *δ*(*x*_*i*_, *e*_*j*_) are also non-negative. Then *C* is the matrix of coefficients of an allocation function because, for every 1 ≤ *j* ≤ *m*, the characterisation from Lemma 2 is satisfied:

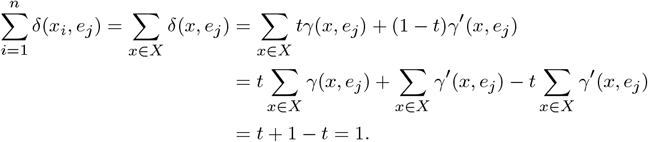

It remains to show that the coefficients *δ*(*x*_*i*_, *e*_*j*_) satisfy the conditions of a diversity index, namely (DI_1_) and (DI_2_). If *x*_*i*_ is not descended from *e*_*j*_, then *γ*(*x*_*i*_, *e*_*j*_) = 0 and *γ* ′ (*x*_*i*_, *e*_*j*_) = 0. Therefore, *δ*(*x*_*i*_, *e*_*j*_) = *t ·* 0 + (1 − *t*) ·0 = 0, and the coefficient *δ*(*x*_*i*_, *e*_*j*_) satisfies (DI_1_).

Next assume that, for some *x, y* ∈ *X* and *e, f* ∈ *E*, to satisfy (DI_2_), we require *γ*(*x, e*) = *γ*(*y, f*). Then *γ* ′ (*x, e*) = *γ* ′ (*y, f*) as well, and hence

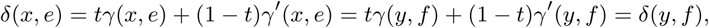

as required. So any set of coefficients that are all equal in *A*_*φ*_ and are also all equal in *B*_*ψ*_ will all be equal in *C*. Thus *C* is the matrix of coefficients of a diversity index, and ***w*** = *C****l*** is contained in *S*(*T, ℓ*). Therefore, *S*(*T, ℓ*) is closed under convex combinations and is a convex space.

□

Suppose that we fix a particular rooted phylogenetic *X*-tree *T* with edge length assignment function *ℓ*. Each possible diversity index for *T* may be viewed as a point inside the convex space *S*(*T, ℓ*), and the Euclidean distances between distinct diversity index vectors indicate their degree of difference. If this distance is zero in *S*(*T, ℓ*), the diversity indices in question coincide on *T* . The space *S*(*T, ℓ*) consists of |*X*| -dimensional vectors. However, it will be shown (Proposition 13) that the vectors in *S*(*T, ℓ*) do not span all of ℝ^*n*^, but rather *S*(*T, ℓ*) ⊂ ℝ^*k*^ for some *k < n*. The smallest such value of *k* is called the *dimension* of *S*(*T, ℓ*). Diversity indices are completely described by their coefficients rather than their edge lengths, so the dimension of *S*(*T, ℓ*) is determined by the tree shape of *T* alone. Although the dimension relies only on the tree shape, the particular boundaries are determined by the edge lengths, as described in Proposition 5.

### 5.1 Examples and the special case when *S*(*T*, .*C*) has dimension zero

To illustrate the effect of tree shape on the dimension of the diversity index space, we examine the spaces of diversity indices for some rooted phylogenetic trees with five leaves. The connection between tree shape and dimension will be formalised in the next section.

Consider again the five-leaf tree in Figure 3. By Proposition 3, the pendant edge lengths are entirely allocated to their incident leaves. By (DI_2_), the length of edge *e*_6_ must be shared equally between *x*_1_ and *x*_2_ in every diversity index. Similarly, the length of *e*_8_ must be shared equally between *x*_4_ and *x*_5_ in every diversity index. Thus, for this tree, only the allocation of edge *e*_7_ between *x*_1_, *x*_2_ and *x*_3_ changes. Suppose that *γ*(*x*_3_, *e*_7_) = *α* is the share of the length of *e*_7_ allocated to *x*_3_. Accordingly, *γ*(*x*_1_, *e*_7_) + *γ*(*x*_2_, *e*_7_) + *α* = 1. Therefore, the share of the length of *e*_7_ allocated to *x*_1_ must equal the share of the length of *e*_7_ allocated to *x*_2_, so 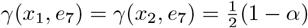. We require 0 ≤ *α* ≤ 1, but *α* is free to be chosen within this range and the stated coefficients will satisfy the diversity index definition. Thus the set of diversity indices on this five-leaf tree form a one-dimensional space, parametrised by the value of *α*. An analogous parametrisation defines the diversity indices on the non-binary tree appearing in Figure 4a.

**Fig. 3:**
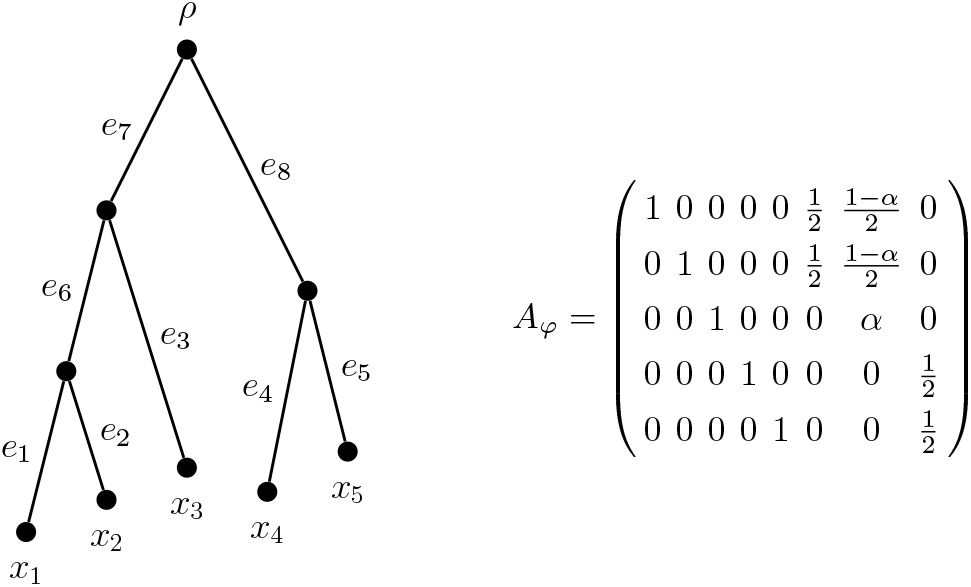
Left: A rooted phylogenetic tree on five leaves. Right: the associated matrix of diversity index coefficients for this tree. The value of 0 ≤ *α* ≤ 1 determines the diversity index (up to coincident indices).

**Fig. 4:**
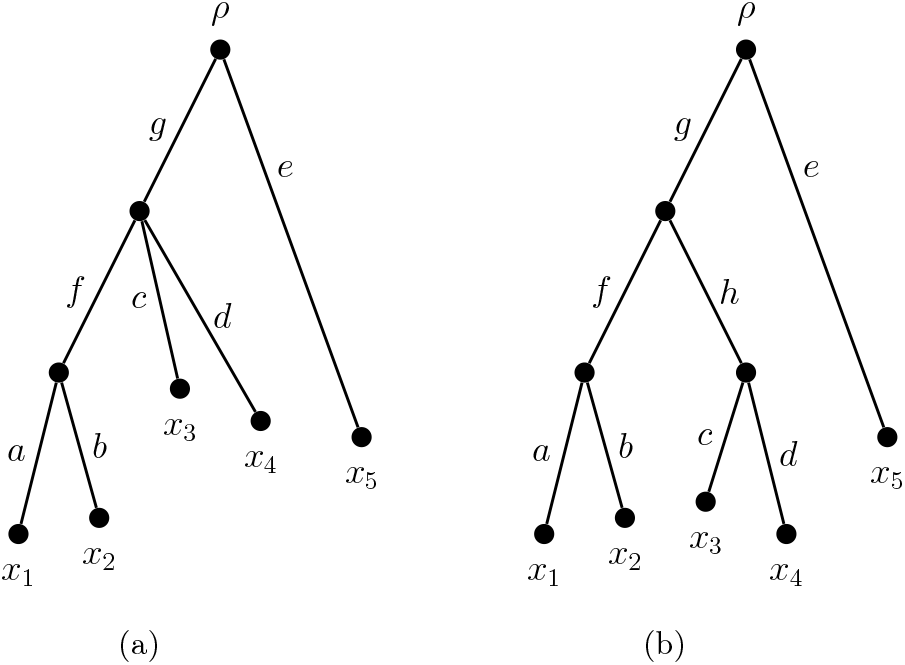
Two rooted phylogenetic tree shapes on five leaves.

For some tree shapes, however, (DI_2_) provides enough of a constraint that all diversity indices coincide. One such shape is given by the tree in Figure 4b, with the necessary index scores appearing in Table 1c. A *balanced* tree is a rooted phylogenetic *X*-tree *T* in which, for every vertex *v*, the pendant subtrees *T*_1_(*v*), …, *T*_*d*_(*v*) all have the same tree shape as each other (where *d* is the outdegree of *v*). We call a pendant subtree *P* of *T maximal* if it is not a subtree of any other pendant subtree, that is, *P* ∈ {*T*_1_(*ρ*), …, *T*_*d*_(*ρ*)}, where *d* is the out-degree of the root vertex *ρ*. A *semi-balanced* tree is a rooted phylogenetic *X*-tree *T* where the maximal pendant subtrees of *T* are all balanced trees.

**Table 1:** Diversity index scores for the trees in Figure 4. Each value of *α*, for 0 ≤ *α* ≤ 1, determines a diversity index for the tree in (a).

| $x$ | $\varphi(x)$ | $x$ | $\varphi(x)$ |
| --- | --- | --- | --- |
| $x_1$ | $\frac{1}{2}(1 - 2\alpha)\ell(g) + \frac{\ell(f)}{2} + \ell(a)$ | $x_1$ | $\frac{\ell(g)}{4} + \frac{\ell(f)}{2} + \ell(a)$ |
| $x_2$ | $\frac{1}{2}(1 - 2\alpha)\ell(g) + \frac{\ell(f)}{2} + \ell(b)$ | $x_2$ | $\frac{\ell(g)}{4} + \frac{\ell(f)}{2} + \ell(b)$ |
| $x_3$ | $\alpha\ell(g) + \ell(c)$ | $x_3$ | $\frac{\ell(g)}{4} + \frac{\ell(h)}{2} + \ell(c)$ |
| $x_4$ | $\alpha\ell(g) + \ell(d)$ | $x_4$ | $\frac{\ell(g)}{4} + \frac{\ell(h)}{2} + \ell(d)$ |
| $x_5$ | $\ell(e)$ | $x_5$ | $\ell(e)$ |

#### Proposition 8

*Let T be a rooted phylogenetic X-tree. The space of diversity indices on T consists of a single point if and only if T is a semi-balanced tree*.

*Proof* Let *φ* (*x*) = Σ_*e*∈*E*(*T*)_ *γ*(*x, e*) *ℓ* (*e*) be a diversity index on *T* . Suppose that *T* is a semi-balanced tree. For every leaf vertex in *T*_1_(*ρ*), there is a symmetry in *T* that swaps that vertex with any other leaf vertex in *T*_1_(*ρ*). A similar argument holds for leaves sharing any other maximal pendant subtree of *T* . Let *e* be an interior edge of *T* . All leaves in *c*_*T*_ (*e*) must belong to the same maximal pendant subtree of *T* . Thus, by Proposition 4, *γ*(*x, e*) = *γ*(*x* ′, *e*) for any *x, x* ′ in *c*_*T*_ (*e*). Therefore, for each leaf *x* descended from *e*, we can set 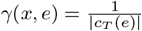. The coefficients associated with every interior edge of *T* can be determined in this way, and those of the pendant edges are fixed by Proposition 3 (iii). Hence, there is only one possible set of index scores, i.e., *S*(*T, ℓ*) has dimension zero for any edge length assignment function *ℓ*.

Conversely, suppose that *T* is not a semi-balanced tree. Then there exists some edge *f* = (*u, v*) (with positive length) such that *T*_1_(*v*) does not have the same tree shape as, say, *T*_2_(*v*). Without loss of generality, assume that there are *a*_1_ maximal pendant subtrees below *v* with the same tree shape as *T*_1_(*v*), and *a*_2_ maximal pendant subtrees below *v* with the same tree shape as *T*_2_(*v*). Let *n*_1_ be the number of leaves in *T*_1_(*v*) and let *n*_2_ be the number of leaves in *T*_2_(*v*). We define the following two distinct diversity indices on *T* .

Firstly, take 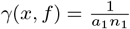 whenever *x* lies in a maximal pendant subtree below *v* that has the same tree shape as *T*_1_(*v*). Then *γ*(*x, f*) = 0 whenever *x* lies in a maximal pendant subtree below *v* that has a different tree shape from *T*_1_(*v*). For a second set of coefficients, take 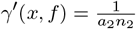 whenever *x* lies in a maximal pendant subtree below *v* that has the same tree shape as *T*_2_(*v*). Then *γ* ′ (*x, f*) = 0 whenever *x* lies in a maximal pendant subtree below *v* that has a different tree shape from *T*_2_(*v*). In both cases, we use the ES coefficients for every other edge (unless an edge has the same descendent structure as *f*, in which case, we copy the pattern of *f* ‘s coefficients in order to satisfy (DI_2_)). Let *φ* (*x*) = Σ_*e ∈%E*_ *γ*(*x, e*) and *ψ*(*x*) =Σ _*e ∈%E*_ *γ*′ (*x, e*). The index scores of *φ* and *ψ* are never coincident, as *ℓ* (*f*) *>* 0. Thus *S*(*T, ℓ*) contains more than one index and cannot consist of a single point.

□

#### Corollary 9

*The Fair Proportion and Equal-Splits indices coincide on a rooted binary phylogenetic tree T if and only if all diversity indices coincide on that tree as well*.

*Proof* Wicke and Steel [7, Theorem. 4] showed that FP and ES coincide on a rooted binary phylogenetic tree *T* if and only if *T* is semi-balanced. Their proof may be extended directly to the non-binary case. By Proposition 8, *T* being semi-balanced is equivalent to *S*(*T, ℓ*) consisting of a single point. That is, all diversity indices on *T* are coincident.

□

## 6 Consistent diversity indices

The coefficients of the FP and ES diversity indices share a property that allows us to view their calculation as a type of flow problem. This is a useful means of understanding the allocation of PD for these indices. For both FP and ES, the allocation of an edge length among the maximal pendant subtrees of a clade is the same for every edge ancestral to that clade, not in the value of the coefficients but in their ratios. For example, consider the tree in Figure 5. The FP allocation of edge *g* to {*x*_1_, *x*_2_} compared with {*x*_3_} is 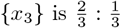, whereas the allocation of edge *h* among these same three species is 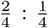. In both cases, a 2:1 ratio of allocations holds. When we use the ES index, the allocation between these same sets follows a 1 : 1 ratio for both *g* and *h*. A diversity index that has the same ratio of allocation at vertex *v*, for every edge that *v* is descended from, we call *consistent at v*. We introduce some notation to aid our discussion of this property. Let 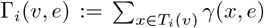, that is, the sum of all coefficients associated with both edge *e* and some leaf in the *i*th pendant subtree below vertex *v*.

**Fig. 5:**
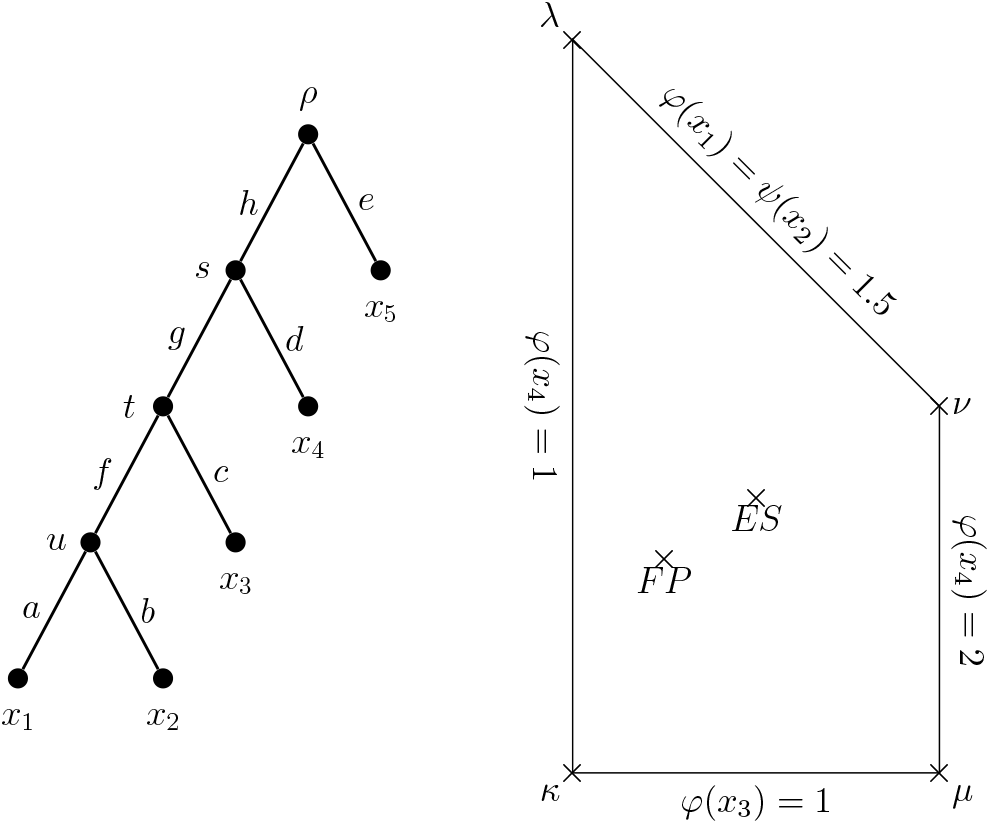
Left: The five-leaf rooted caterpillar tree *Cat*_5_, where each edge has unit length. Right: The diversity index space *S*(*Cat*_5_, **1**), projected onto the *φ*(*x*_3_) *φ*(*x*_4_)-plane, with boundary conditions and extremal indices *κ, λ, μ, ν* at the corner points. Notice that the FP and ES diversity indices appear as interior points of this convex space.

### Definition 5

(Consistency condition) Let *T* = (*V, E*) be a rooted phylogenetic *X*tree. Choose *e, f* ∈ *E*, with *e* = (*u, v*) descended from *f*, and let *T*_1_(*e*), …, *T*_*d*_(*e*) be the maximal pendant subtrees descended from the terminal vertex of *e*. A diversity index *φ* (*x*) = Σ_*e*∈*E*_ *γ*(*x, e*) *ℓ* (*e*) is *consistent at v* if there exists a constant *k* ∈ ℝ^≥0^ such that *Γ*_*i*_(*v, f*) = *kΓ*_*i*_(*v, e*). If *φ* is consistent at *v* for every *v* ∈ *V*, then we say that *φ* itself is *consistent*.

At vertex *v*, descended from *e ∈ E*, the ratio *Γ*_1_(*v, e*) : … : *Γ*_*d*_(*v, e*) is called the *ratio of allocations* at *v*. A ratio of allocations is *normalised* if it has been scaled by some *t* ∈ ℝ such that 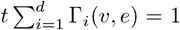. Reiterating our earlier example, a consistent diversity index on the rooted phylogenetic tree in Figure 5 requires the ratio of allocation [*γ*(*x*_1_, *g*) + *γ*(*x*_2_, *g*)] : *γ*(*x*_3_, *g*) to be the same as [*γ*(*x*_1_, *h*) + *γ*(*x*_2_, *h*)] : *γ*(*x*_3_, *h*).

Consistent diversity indices have the convenient property that their index scores are able to be determined by a simple flow-based algorithm. The idea is to view each index as a rule for re-weighting edges, moving weight from each interior edge to its immediate descendent edges. The algorithm begins with the edges incident to the root vertex and continues to re-weight edges until all interior edges have a weight of zero. The diversity index score of each leaf is then given by the final length of its incident pendant edge. This approach is reminiscent of the transformations of edge lengths in [19] that maintain the Shapley values of each leaf.

Consider a particular rooted phylogenetic tree *T* with fixed edge lengths *ℓ*. We now show that any diversity index on *T* may be framed as a consistent index. This reframing, in turn, will allow us to describe the dimension and extreme points of *S*(*T, ℓ*).

### Lemma 10

*Let T* = (*V, E*) *be a rooted phylogenetic X-tree with a fixed edge length assignment function ℓ. Let φ* (*x*) = Σ_*e ∈%E*_ *γ*(*x, e*) *ℓ* (*e*) *be a consistent diversity index on T* . *Given the ratios of allocation at each vertex, we can reconstruct the diversity index coefficients*.

*Proof* For each *v* ∈ *V*, we normalise the ratios of allocation, writing the scaled ratio as *r*_1_(*v*) : *…* : *r*_*d*_(*v*). Let *e* = (*u*_0_, *u*_1_) be an interior edge of *T* and let *x* ∈ *X* be descended from *e*. Let *P* (*T* ; *u*_1_, *x*) consist of the vertices {*u*_1_, *u*_2_, …, *u*_*k*_, *x*} and without loss of generality assume that *x* ∈ *X* is in *T*_1_(*v*) for all *v* ∈ *P* (*T* ; *u*_1_, *u*_*k*_).

Firstly, the proportion of *ℓ* (*e*) allocated among vertices in *T*_1_(*u*_1_) is *r*_1_(*u*_1_). Next note that the proportion of *ℓ* (*e*) allocated among vertices in *T*_1_(*u*_2_) is *r*_1_(*u*_2_) of the *ℓ* (*e*) allocation coming into *u*_2_. That is, *r*_1_(*u*_1_) · *r*_1_(*u*_2_) overall. Similarly, the leaves of *T*_1_(*u*_3_) are allocated a total of *r*_1_(*u*_1_) · *r*_1_(*u*_2_) · *r*_1_(*u*_3_) of *ℓ* (*e*). Continuing in this manner until the parent of *x*, we find that *γ*(*x, e*) is given by the product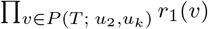. For an arbitrary leaf ′ descended from edge *e* ′, the coefficient *γ*(*x* ′, *e* ′) can be expressed as a similar product, taking the appropriate ratio terms at vertices along the path connecting edge *e* ′ to leaf *x* ′.

□

We give a short example of the calculation above in order to clarify the final sentence of this proof. Suppose that *x* is a leaf of a phylogenetic tree rooted at *ρ*, and that the path from *ρ* to *x* in *T* passes through vertices *s, t, u* and *v* in turn. Let *e* be the edge (*s, t*). Suppose further that *x* ∈ *T*_2_(*t*), *x* ∈ *T*_5_(*u*) and *x* ∈ *T*_1_(*v*). Then the coefficient *γ*(*x, e*) in this case is given by the product *r*_2_(*t*) · *r*_5_(*u*) · *r*_1_(*v*).

### Proposition 11

*Let T* = (*V, E*) *be a rooted phylogenetic X-tree, rooted at ρ, with a fixed edge length assignment function £. Let φ* (*x*) = Σ_*e*∈*E*_ *γ*(*x, e*) *ℓ* (*e*) *be a diversity index on T* . *Then there exists a consistent diversity index ψ*_*ℓ*_ (*x*) = Σ_*e*∈*E*_ *γ* ′ (*x, e*) *ℓ* (*e*) *such that φ and ψ*_*ℓ*_ *coincide*.

*Proof* We construct a consistent diversity index *ψ*_*ℓ*_ (*x*) = Σ_*e ∈%E*_ *γ*′ (*x, e*) *ℓ* (*e*) in the following manner. Let *v* be a vertex of *T* with out-degree *d*. We take the ratio of allocations for *ψ*_*ℓ*_ at *v*, using the coefficients from *φ* and the edge lengths given by *ℓ*, as

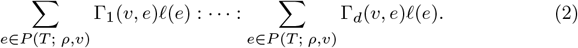

Let *r*_1_(*v*) : *…* : *r*_*d*_(*v*) be the normalised ratio equivalent to ratio 2. We construct the normalised ratio of allocations at each interior non-root vertex of *V* in a similar way. Then applying the method of Lemma 10 gives the *γ′* (*x, e*) coefficients of *ψ*_*ℓ*_.

We now show that *ψ*_*ℓ*_ coincides with *φ* for an arbitrary leaf *x* ∈ *X*. By the descent condition (DI_1_), we need only consider edges from the path 𝒫 = *P* (*T* ; *ρ, x*). Specifically, let 𝒫 consist of the vertices {*ρ, u*_1_, *u*_2_, …, *u*_*k*_, *x*} and edges *e*_1_ = (*ρ, u*_1_), *e*_2_ = (*u*_1_, *u*_2_), …, *e*_*k*+1_ = (*u*_*k*_, *x*). Assume, without loss of generality, that *x* is in *T*_1_(*v*) for all *v* ∈ 𝒫 \ {*ρ, x*}.

Using ratio 2 and normalising gives values of 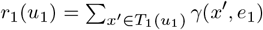 and 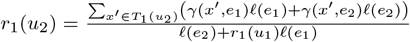 and so on until *T*_1_ (*u*_*k*_) contains only the leaf *x*, where

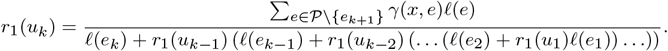

By Lemma 10, the coefficients of *ψ*_*ℓ*_ are given by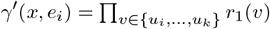. So

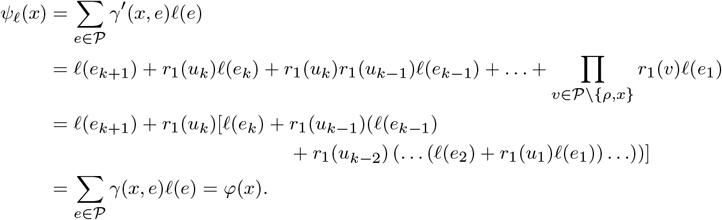

Therefore, as *x* was arbitrary, the consistent diversity index *ψ*_*ℓ*_ coincides with *φ*.

□

Let *T* = (*V, E*) be a rooted phylogenetic *X* -tree with fixed edge length assignment function *ℓ*. To each interior non-root vertex *v* we associate an integer: the *degrees of freedom* of *v*. This value is calculated as one less than the number of distinct tree shapes across the set of maximal pendant subtrees below *v* (i.e. in *T*_1_(*v*), …, *T*_*d*_(*v*)). In a rooted binary phylogenetic tree, each interior non-root vertex therefore has zero or one degree of freedom.

We construct an equivalence relation ∼ on the set of interior non-root vertices of *T* . We write *u* ∼ *v* if and only if *u* and *v* both have out-degree *d*, and the multiset of tree shapes of {*T*_1_(*u*), …, *T*_*d*_(*u*)} equals the multiset of tree shapes of *T*_1_(*v*), …, *T*_*d*_(*v*) . In other words, *u* ∼ *v* if and only if the structure of the subtree descended from *u* is exactly the same as that descended from *v*. Theorem 12 allows us to use the ∼ -equivalence classes to determine the dimension of each diversity index space exactly.

### Theorem 12

*Let T* = (*V, E*) *be a rooted phylogenetic X-tree with a fixed edge length assignment function ℓ. Let V* ′ *be a set that contains precisely one vertex from each of the ~-equivalence classes of T* . *The dimension of the convex space S*(*T, ℓ*) *of diversity indices is the sum of the degrees of freedom of the vertices in V* ′.

*Proof* For each *v* ∈ *V* ′, consider the normalised ratio of allocations *r*_1_(*v*) : *…* : *r*_*d*_(*v*) (where *d* is the out-degree of *v*). Assume that *v* is an interior non-root vertex with zero degrees of freedom. Then to satisfy condition (DI_2_) requires that 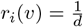 for all *i* ∈ {1, …, *d*}. Next, assume that *v* is an interior non-root vertex with non-zero degrees of freedom. Suppose that there are *k*_1_ maximal pendant subtrees with the same tree shape as *T*_1_(*v*). Any value in the interval 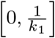 is possible for *r*_1_(*v*). If *T*_*i*_(*v*) has the same tree shape as *T*_1_(*v*), then set *r*_*i*_(*v*) equal to *r*_1_(*v*).

As *v* has non-zero degrees of freedom, there is a maximal pendant subtree, say *T*_2_(*v*), with a tree shape that is different from *T*_1_(*v*). Suppose that there are *k*_2_ maximal pendant subtrees with the same tree shape as *T*_2_(*v*). If *v* has exactly one degree of freedom, then 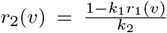. Otherwise, any value in the interval 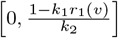 is possible for *r*_2_(*v*), and those other terms of the ratio corresponding to a tree shape matching *T*_2_(*v*). This process continues until the number of values selected matches the degrees of freedom of *v*. Then the ratio term(s) corresponding to the last tree shape (among the maximal pendant subtrees of *v*) is the value which ensures that the ratio of allocations is normalised. Finally, if *u* ∼ *v* and *T*_*i*_(*u*) has the same tree shape as *T*_*j*_ (*v*), we set *r*_*i*_(*u*) = *r*_*j*_ (*v*).

For each ∼-equivalence class of interior non-root vertices with non-zero degrees of freedom, we can choose the ratio terms independently. We now form a vector where each component corresponds to a choice of a ratio term by the process above. There is thus one component per degree of freedom (in total, across all equivalence classes). By Lemma 10, for each possible set of ratios there is a unique set of coefficients and thus a unique corresponding diversity index. Hence, the dimension of *S*(*T, ℓ*) is at least the sum of the degrees of freedom of the vertices in *V* ′.

On the other hand, Proposition 11 shows that any diversity index in *S*(*T, ℓ*), say *φ*, coincides with a consistent diversity index, say 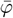. At each vertex, the terms in the normalised ratio of 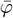 must lie within the intervals described in our construction above. Otherwise, the sum of the ratio terms would add to more than one, or else some of the ratio terms would be negative. Both possibilities are excluded by the definition of a normalised ratio of allocations. Thus it is possible to construct 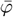 in the manner described above. Hence, the dimension of the convex space *S*(*T, ℓ*) is precisely the sum of the degrees of freedom of the vertices in *V* ′. □

We note that the sum of degrees of freedom will always be less than the number of leaves in a given rooted phylogenetic tree and express this inequality through the following result.

### Proposition 13

*Let T be a rooted phylogenetic X-tree with fixed edge lengths £. The dimension of S*(*T, ℓ*) *is at most* |*X*| − 2.

*Proof* The dimension of *S*(*T, ℓ*) is calculated by adding up the degrees of freedom in each ∼-equivalence class. We show that even when adding together degrees of freedom from every interior non-root vertex (as opposed to just one per ∼-equivalence class), that the sum of degrees of freedom always remains at most |*X*| − 2.

Let *n* = |*X*| and let *df* (*T*) be the total degrees of freedom summed across every interior non-root vertex in *T* . Then *df* (*T*) is at least the dimension of *S*(*T, ℓ*). Furthermore, let I(*T*) be the number of interior non-root vertices of *T* and let *i*(*d*) be the number of interior vertices of *T* that have out-degree *d*. We consider every combination of interior vertices, in terms of their out-degrees. The total *df* (*T*) value is comprised of, at most, one degree of freedom per an extra degree of freedom for each third and subsequent edge originating from an interior non-root vertex. In symbols, 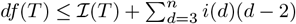.

The number of interior non-root vertices of *T* can be determined by starting with an appropriate binary *X*-tree *T* ′ and contracting edges to form *T* . The number of such contractions required is *d* − 2 for each interior vertex with degree *d* ≥ 3 in *T* and each reduces the number of interior vertices by one. As every binary *X*-tree has *n* − 2 interior non-root vertices, 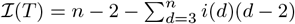.

Finally, this gives 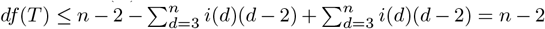.Therefore the total number of degrees of freedom of *T*, and hence *S*(*T, ℓ*), is at most |*X*| − 2. □

### 6.1 Example: application to a tree of hominoids

A brief illustration of this is provided by the rooted phylogenetic tree of Hominoids appearing in Figure 6. For this tree, each of the nine labelled interior vertices is a representative of a ∼ -equivalence class that has one degree of freedom. There are also two equivalence classes among the unfilled circle vertices, each with zero degrees of freedom. Hence, there are nine degrees of freedom overall for this tree, and the diversity index space for the tree of hominoids has nine dimensions. In other words, any diversity index on this tree is required to specify the ratio of allocation at these nine labelled representative vertices in order to determine its full set of coefficients.

**Fig. 6:**
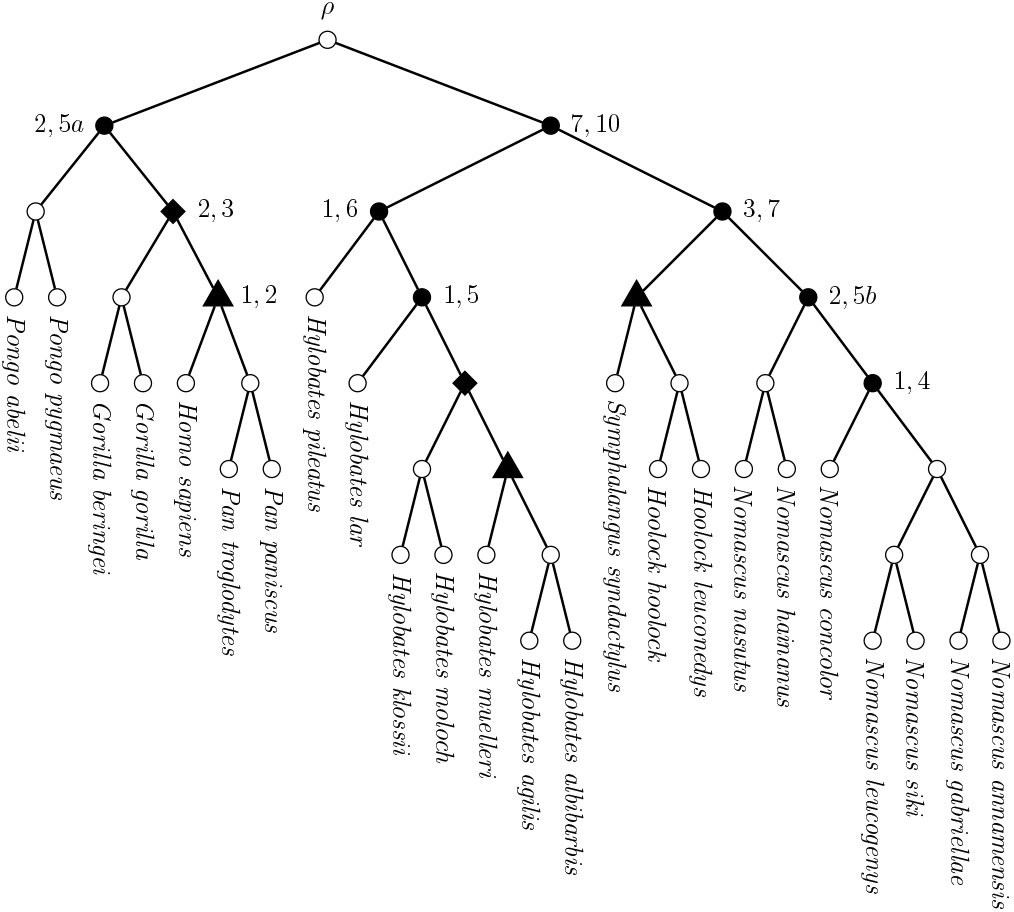
The phylogenetic tree of the superfamily of Hominoids (great apes and gibbons). This tree was constructed using data from [20, 21], via [22]. Filled vertices represent interior vertices that have one degree of freedom. Each numerical label indicates a representative of each ∼ -equivalence class. The two diamond-shaped vertices both belong to the 2,3 class, and the three triangular vertices belong to the 1,2 class. Note that the vertices labelled 2,5a and 2,5b lie in distinct ∼ -equivalence classes because of the difference in tree shape between their maximal pendant subtrees with five leaves. Thus, there are nine equivalence classes, contributing one degree of freedom each.

**Fig. 7:**
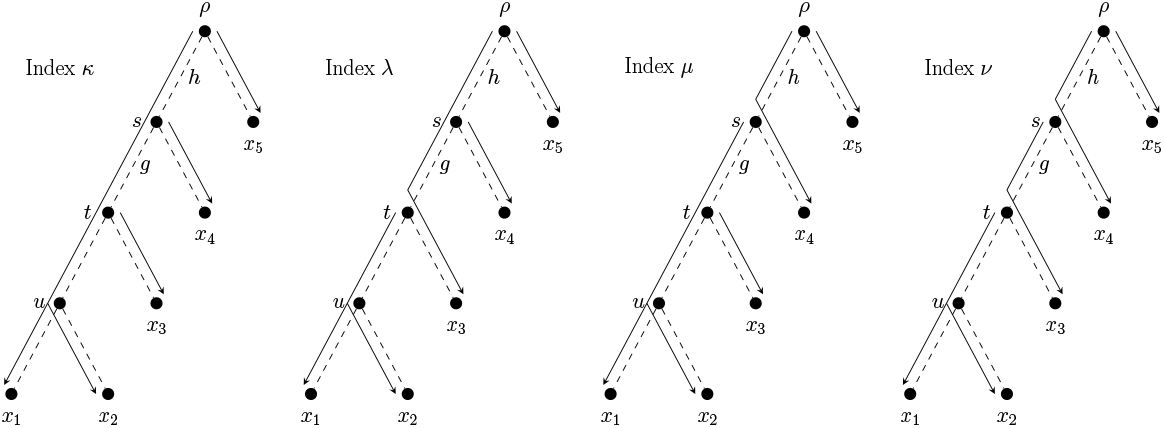
Arrows show the allocations for the four diversity indices on *Cat*_5_ (indicated by dashed edges) that lie at the corners of *S*(*Cat*_5_, **1**).

The presence of a nine-dimensional diversity index space here does not particularly illuminate anything about hominoid-specific diversity. However it does illustrate the point that for even quite a modestly-sized phylogenetic tree as this, a large number of potential diversity indices exist. Moreover, most potential diversity indices that could be applied to the hominoid tree are not directly related to the known FP and ES indices.

The implication for conservation is that more investigation of the diversity index space is required to ensure optimal use of the diversity index concept. This would involve testing various properties of candidate diversity indices against the assumption that FP or ES is the most suitable index (from among the infinitely many possibilities). Doing so could help establish whether FP, ES or some new diversity index (or indices) best give a meaningful and robust measurement of a taxon’s phylogenetic isolation and embodiment of shared and unique evolutionary history.

### 6.2 Boundaries of index spaces

We give an example of a two-dimensional diversity index space on the rooted caterpillar tree on five leaves (Figure 5), which we denote here as *Cat*_5_. Let *φ* be a diversity index on *Cat*_5_, and suppose that every edge in *Cat*_5_ has unit length; we denote this as ***ℓ***= **1**. By taking the extreme ratios of allocation at vertices *s* and *t* (see Table 2), we find the diversity indices *κ, λ, μ* and *ν* (index score vectors listed below) that lie at the boundary points of *S*(*Cat*_5_, **1**):

**Table 2:** The corners of *S*(*Cat*_5_, **1**) are determined by taking the four possible combinations of extreme ratios at *s* and *t*.

| Index | Ratio at $s$ | Ratio at $t$ |
| --- | --- | --- |
| $\kappa$ | 1:0 | 1:0 |
| $\lambda$ | 1:0 | 0:1 |
| $\mu$ | 0:1 | 1:0 |
| $\nu$ | 0:1 | 0:1 |

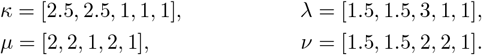

Similarly, the ‘corner’ indices of a diversity index space may be found by taking the extreme ratios of allocation for each ∼ -equivalence class in the corresponding phylogenetic tree. We can then use these corner indices to obtain further diversity indices by way of linear combination. Carathéodory’s Theorem (this version was drawn from Steinitz [23]) describes how each point of a convex space may be described as a combination of a limited number of points.

#### Theorem 14

(Carathéodory’s Theorem) *Let A be a non-empty subset of* ℝ^*d*^. *Every vector from the convex hull of A can be represented as a convex combination of, at most, d* + 1 *vectors from A*.

Specifically, for a rooted phylogenetic tree *T*, we can construct the required *d* + 1 vectors as follows. If *T* is semi-balanced, then only one vector is required, so we can choose that belonging to FP. Otherwise, for each ∼-equivalence class, we choose an extreme ratio of allocation and let the resulting diversity index be called *B*_1_. Next, we construct a new diversity index *B*_2_ by matching the ratios of allocation from *B*_1_, except that for precisely one of the ∼ -equivalence classes, an alternative extreme ratio of allocation is chosen. Exactly *d* such diversity indices may be constructed in addition to *B*_1_, one for each degree of freedom in the tree. The resulting diversity indices *B*_1_, …, *B*_*d*+1_ cover the total degrees of freedom of *T* . Hence *B*_1_, …, *B*_*d*+1_ are the required *d* + 1 diversity indices, from which all others in *S*(*T, ℓ*) may be constructed.

Continuing our example, *S*(*Cat*_5_, **1**) ⊂ ℝ^2^ is itself convex. Therefore the vector of index scores of any diversity index in this space may be expressed as a convex combination of at most three points of *S*(*Cat*_5_, **1**). Specifically, we can express any diversity index *φ* for *Cat*_5_ as

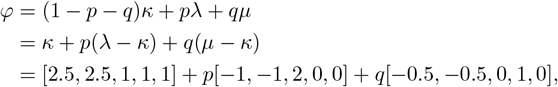

where 0 ≤ *p, q* ≤ 1, provided that 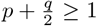. FP is given by 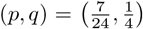, and ES is given by 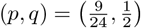. These positions are noted in Figure 5.

## 7 Concluding remarks

In this paper we have investigated the combinatorial and geometric properties of the space of phylogenetic diversity indices, having defined these in a very general way. The benefit of doing so is that one can more readily investigate which properties of known diversity indices hold generally and which are specific to a restricted class of diversity indices. Understanding the properties of diversity indices may be useful when deciding which index to use in a particular setting. For example, we have discussed the continuity property of the Fair Proportion index and shown that it is unique to that index.

We briefly present two further properties that may be of interest in this regard. Each are held by both the FP and ES indices. Let *T* = (*V, E*) be a rooted phylogenetic *X*-tree, and let *e* = (*u, v*) ∈ *E*.

- Property 1: the ratio of allocations *Γ*_1_(*v, e*) : *…* : *Γ*_*d*_(*v, e*) depends only on the number of leaves in each of the *d* maximal pendant subtrees below *e*, and not their structure.
- Property 2: *Γ*_*i*_(*v, e*) ≥ *Γ*_*j*_(*v, e*) whenever *T*_*i*_(*v*) contains at least as many leaves as *T*_*j*_(*v*).

We have focussed on a particular structure, namely that of rooted phylogenetic trees. For consistent diversity indices, the ability to view their calculation as a flow problem allows a straightforward extension of this framework to rooted phylogenetic networks. In the more general setting of a phylogenetic network, we need only to add the stipulation that the flow into a so-called reticulation vertex is matched by the flow out from that vertex. A second more general approach could be developed by applying allocation functions to unrooted phylogenetic trees. It is for this reason that we have presented the definition of a diversity index as a subclass of allocation functions, although investigation of allocating PD among leaves of unrooted trees has been left for further work.

## Acknowledgments

The authors were supported by the New Zealand Marsden Fund (MFP-UOC2005). We thank Martin Frohn for useful suggestions regarding both Proposition 5 and specifying a ‘basis’ for diversity index spaces.

## Declarations

The authors have no relevant financial or non-financial interests to disclose. The authors have no conflicts of interest to declare that are relevant to the content of this article. The authors certify that they have no affiliations with or involvement in any organisation or entity with any financial interest or non-financial interest in the subject matter or materials discussed in this manuscript. The authors have no financial or proprietary interests in any material discussed in this article.

## Footnotes

1 Alternative ranking approaches, such as the methods of [9] or [10], may be useful when the structure of a phylogeny is known but its edge lengths are uncertain. However, we do not consider these further here.

## Notes

### Competing Interest Statement

The authors have declared no competing interest.

### Summary of Updates

More details provided for Lemma 10, Proposition 11. New direct proof of Proposition 13 in place of earlier proof by contradiction and other minor changes to improve clarity of presentation throughout.

## References

[1] Felsenstein, J.: Inferring Phylogenies. Sinauer Associates, Sunderland, MA, USA (2004)

[2] Cadotte, M.W., Jonathan Davies, T.: Rarest of the rare: advances in combining evolutionary distinctiveness and scarcity to inform conservation at biogeographical scales. Diversity and Distributions 16(3), 376–385 (2010)

[3] Tucker, C.M., Cadotte, M.W., Carvalho, S.B., Davies, T.J., Ferrier, S., Fritz, S.A., Grenyer, R., Helmus, M.R., Jin, L.S., Mooers, A.O., et al.: A guide to phylogenetic metrics for conservation, community ecology and macroecology. Biological Reviews 92(2), 698–715 (2017)

[4] Faith, D.P.: Conservation evaluation and phylogenetic diversity. Biological Conservation 61, 1–10 (1992)

[5] Hartmann, K.: The equivalence of two phylogenetic biodiversity measures: the Shapley value and fair proportion index. Journal of Mathematical Biology 67(5), 1163–1170 (2013)

[6] Redding, D.W., Mazel, F., Mooers, A.Ø.: Measuring evolutionary isolation for conservation. PLoS One 9(12), 113490 (2014)

[7] Wicke, K., Steel, M.: Combinatorial properties of phylogenetic diversity indices. Journal of Mathematical Biology 80(3), 687–715 (2020)

[8] Fischer, M., Francis, A., Wicke, K.: Phylogenetic diversity rankings in the face of extinctions: The robustness of the fair proportion index. Systematic Biology (2022)

[9] Crozier, R.H.: Genetic diversity and the agony of choice. Biological conservation 61(1), 11–15 (1992)

[10] Vane-Wright, R.I., Humphries, C.J., Williams, P.H.: What to protect?—systematics and the agony of choice. Biological conservation 55(3), 235–254 (1991)

[11] EDGE of Existence Programme: EDGE of Existence: Evolutionarily Distinct & Globally Endangered (2022). http://www.edgeofexistence.org Accessed 3 July 2022

[12] Gumbs, R., Gray, C.L., Böhm, M., Burfield, I.J., Couchman, O.R., Faith, D.P., Forest, F., Hoffmann, M., Isaac, N.J.B., Jetz, W., et al.: The EDGE2 protocol: advancing the prioritisation of evolutionarily distinct and globally endangered species for practical conservation action. PLoS Biology (In Press)

[13] Isaac, N.J., Turvey, S.T., Collen, B., Waterman, C., Baillie, J.E.: Mammals on the edge: conservation priorities based on threat and phylogeny. PloS One 2(3), 296 (2007)

[14] Palmer, C., Fischer, B.: Should global conservation initiatives prioritize phylogenetic diversity? Philosophia, 1–20 (2021)

[15] Redding, D.: Incorporating genetic distinctness and reserve occupancy into a conservation priorisation approach. Master’s thesis: University of East Anglia (2003)

[16] Wicke, K.: Novel aspects of mathematical phylogenetics. PhD thesis, Universität Greifswald (2020)

[17] Steel, M.: Phylogeny: Discrete and Random Processes in Evolution. SIAM, Philadelphia, PA, USA (2016)

[18] Fuchs, M., Jin, E.Y.: Equality of shapley value and fair proportion index in phylogenetic trees. Journal of mathematical biology 71, 1133–1147 (2015)

[19] Haake, C.-J., Kashiwada, A., Su, F.E.: The Shapley value of phylogenetic trees. Journal of Mathematical Biology 56(4), 479–497 (2008)

[20] Springer, M.S., Meredith, R.W., Gatesy, J., Emerling, C.A., Park, J., Rabosky, D.L., Stadler, T., Steiner, C., Ryder, O.A., Janečka, J.E., et al.: Macroevolutionary dynamics and historical biogeography of primate diversification inferred from a species supermatrix. PloS One 7(11), 49521 (2012)

[21] Carbone, L., Alan Harris, R., Gnerre, S., Veeramah, K.R., Lorente-Galdos, B., Huddleston, J., Meyer, T.J., Herrero, J., Roos, C., Aken, B., et al.: Gibbon genome and the fast karyotype evolution of small apes. Nature 513(7517), 195–201 (2014)

[22] OneZoom Core Team: OneZoom Tree of Life Explorer Version 3.5 (2021). http://www.onezoom.org Accessed 3 July 2022

[23] Steinitz, E.: Bedingt konvergente Reihen und konvexe Systeme. Journal für die reine und angewandte Mathematik 144, 1–40 (1914)

